# Tiled Reconstruction Improves Structured Illumination Microscopy

**DOI:** 10.1101/2020.01.06.895318

**Authors:** David P. Hoffman, Eric Betzig

**Author notes:** Eikon Therapeutics, Inc. Hayward, CA 94545, USA.

## Abstract

Structured illumination microscopy (SIM) is widely used for fast, long-term, live-cell super-resolution imaging. However, SIM images can contain substantial artifacts if the sample does not conform to the underlying assumptions of the reconstruction algorithm. Here we describe a simple, easy to implement, process that can be combined with any reconstruction algorithm to alleviate many common SIM reconstruction artifacts and briefly discuss possible extensions.

## Main Text

Among super-resolution methods, structured illumination microscopy (SR-SIM)^1^ stands out for its ability to image live cells in multiple colors and at high speed with minimal phototoxicity over large fields of view (FOV).^2–4^ However, one criticism often leveled against SR-SIM^5^ is that its reconstruction algorithm employs both experimental and user-defined parameters which, if inaccurately measured or poorly chosen, can lead to image artifacts.^6^ For example, even measurement errors as small as 0.2% in SR-SIM illumination parameters (direction, phase, period, and modulation amplitude) have been shown to lead to substantial artifacts.^3,6^

Traditional SR-SIM^7^ assumes that the reconstruction parameters remain constant over the entire FOV. However, in practice optical aberrations can locally perturb illumination parameters and the detection point spread function (PSF) while differences in thickness or volumetric fluorescence density within the sample can lead to spatial variations in the optimal values of user-defined parameters for noise filtering, apodization, and discrete spatial frequency suppression needed to minimize artifacts.^6^ Spatial variation of the PSF is also commonly encountered in deconvolution^8^ where one stratagem to minimize artifacts has been to divide the FOV into subregions over which the PSF is assumed or known to be nearly constant.^9,10^ Here we apply tiled reconstruction, which has been used to reduce stripe artifacts in optical sectioning SIM,^11^ to SR-SIM by dividing each raw SR-SIM data set into overlapping tiled subsets. Each subset can be reconstructed with independently measured or user-optimized parameters and then reassembled into a final, blended SR-SIM image covering the original FOV (bottom of Supplementary Fig. 1).

To illustrate, we collected cryogenic SR-SIM^12^ data of a high pressure frozen cell expressing the endoplasmic reticulum (ER) lumen marker mEmerald-ER3, and compared standard (Fig. 1a, top) and tiled (Fig. 1a, bottom) reconstruction. In many regions the reconstructed images were similar (Fig. 1b), but in others conventional reconstruction yielded artifacts not apparent with the tiled approach. Spatial maps of the illumination parameters across all tiles revealed particularly large variations in the standing wave modulation amplitude (Supplementary Fig. 2, right column) which, when approximated with a single global value in conventional reconstruction, could lead to such artifacts.

**Fig. 1:**
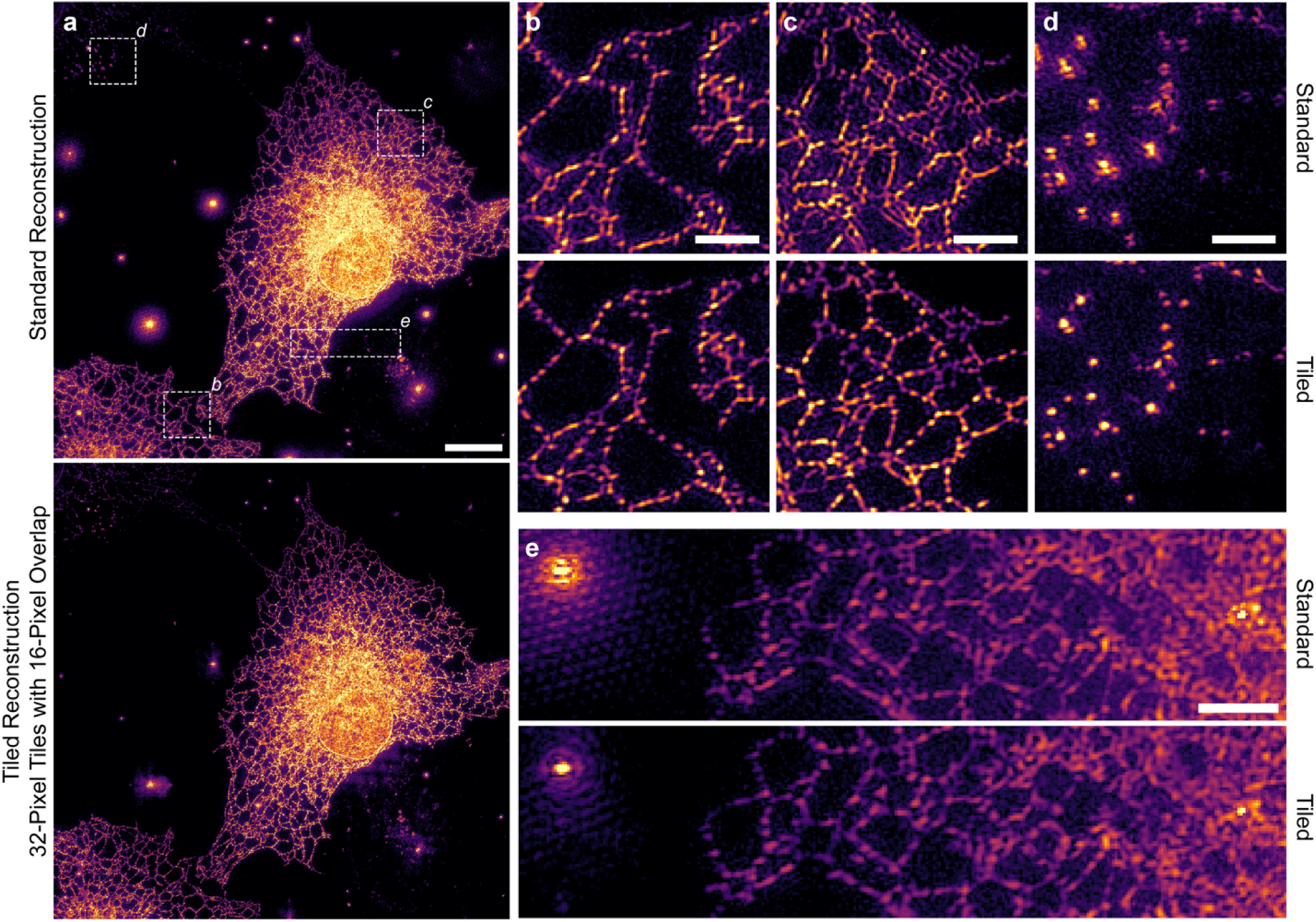
Tiled SR-SIM reconstruction minimizes common SR-SIM artifacts. (**a**, top) Maximum intensity projection (MIP) through a standard 3D-SR-SIM reconstruction of a high pressure frozen COS-7 cell expressing an ER luminal marker. (**a**, bottom) Tiled reconstruction of the same cell using 1024 tiles, each 32 × 32-pixels with 16-pixel overlap. Scalebar, 10 *μ*m. (**b**) An area of the cell with accurate reconstruction by both standard (top) and tiled (bottom) approaches. (**c**) An area of the cell exhibiting a “ghosting” artifact, under standard reconstruction (top) that is not present under tiled reconstruction (bottom). (**d**) Elimination of ringing artifacts at puncta through tiled reconstruction. (**e**) Crosshatch artifacts common in bright structures are effectively suppressed by tiled reconstruction (bottom). Colormaps are scaled identically for each pair of images. Scalebars, 2 *μ*m.

Tiled reconstruction introduces a new parameter, tile size. While smaller tile sizes ensure less variation in the reconstruction parameters across each tile, there is a practical limit below which the effects of pixilation become large and degrade the resulting images (right column of Supplementary Fig. 3). This occurs when, in Fourier space, the number of pixels in the overlap region between the diffraction limited information region and the frequency shifted information regions containing super-resolution spatial frequencies become too small to be stitched together accurately (Supplementary Fig. 4). Although we found that even 16-pixel tiles can produce accurate reconstructions (column 5 of Supplementary Fig. 3), in practice 64-pixel tiles offer a good compromise between minimizing artifacts from spatial variations in reconstruction parameters and maximizing reconstruction robustness and computational efficiency.

We can envision a few extensions to tiled reconstruction. First, thicker specimens might benefit from tiling in 3D rather than only laterally as shown here. Second, square tile shapes are arbitrary, and fewer tiles might be needed if chosen on the basis of sample characteristics leading to spatially varying reconstruction parameters, such as sample thickness, proximity to the nucleus, or local fluorescence density. Third, if sample-induced aberrations affecting the detection pathway spatially vary, one could use a locally measured PSF for each tile during reconstruction. Regardless, we have found that tiled reconstruction is a simple and effective expedient for reducing artifacts in SR-SIM images that should benefit the broader SR-SIM community.

## Acknowledgements

We thank M. Freeman for preparing the samples, G. Shtengel for help in constructing the microscope, and L. Shao and D. Li for SR-SIM assistance. mEmerald-ER3 were a gifts from Michael Davidson (Addgene plasmid #54082; http://n2t.net/addgene:54082; RRID:Addgene_54082).

## Contributions

D.P.H. conceived the concept, collected and processed the data. E.B. supervised the project and wrote the correspondence together with D.P.H.

## Ethics declarations

### Competing interests

The authors declare no competing interests.

