## Supplemental Material for "Tiled Reconstruction Improves Structured Illumination Microscopy"

### Supplementary Information

#### Supplementary Methods

The SR-SIM microscope used here[12] is similar to one described previously[2] except the specimen holder is located within an optical cryostat in order to image vitreously frozen specimens. Cells were frozen using a high pressure freezer (Wohlwend Compact 2, Technotrade International) after transfection and plating on custom sapphire coverslips (3 mm diameter, 0.05 mm thickness, Nanjing Co-Energy Optical Crystal Co., Ltd).

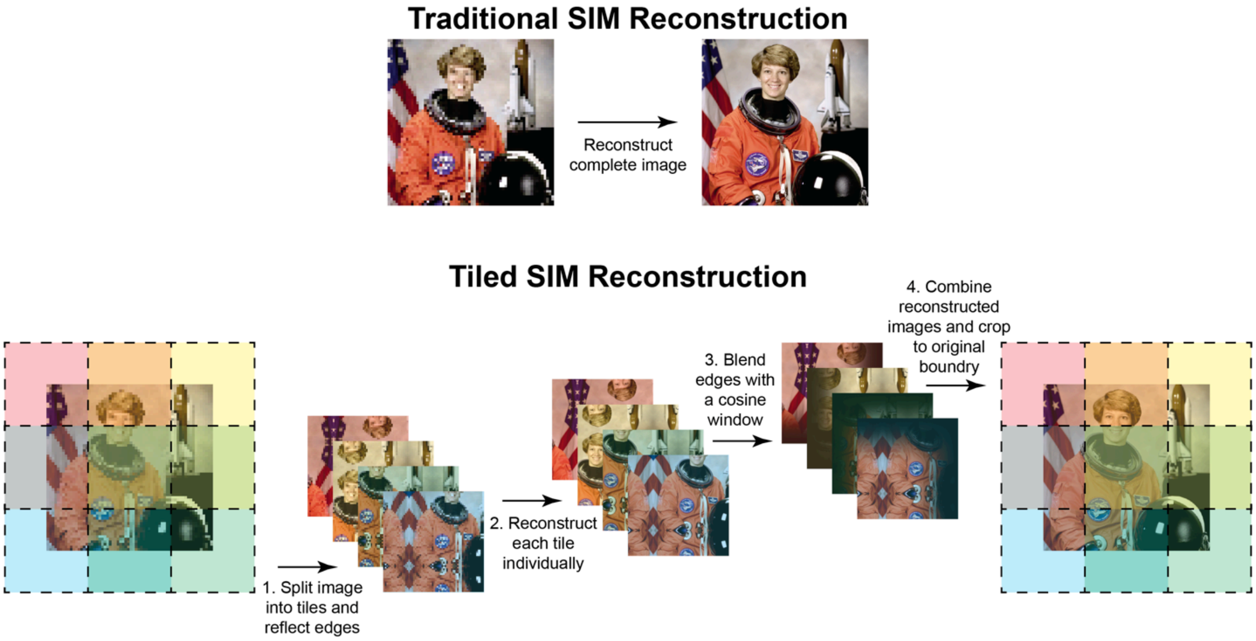


**Supplementary Fig. 1.** Tiled Reconstruction Algorithm.

Traditional SR-SIM (top) applies a single set of reconstruction parameters across the entire field of view (FOV).  Tiled SR-SIM reconstruction (bottom) breaks the image data into smaller image tiles and reconstructs them independently. Tiles have overlapping edges which are blended with their neighbors using a cosine window to assemble the final reconstructed image. Pixels in the reconstructed examples are shown 8X smaller than the raw images for illustrative purposes, rather than the true 2X difference in SR-SIM,

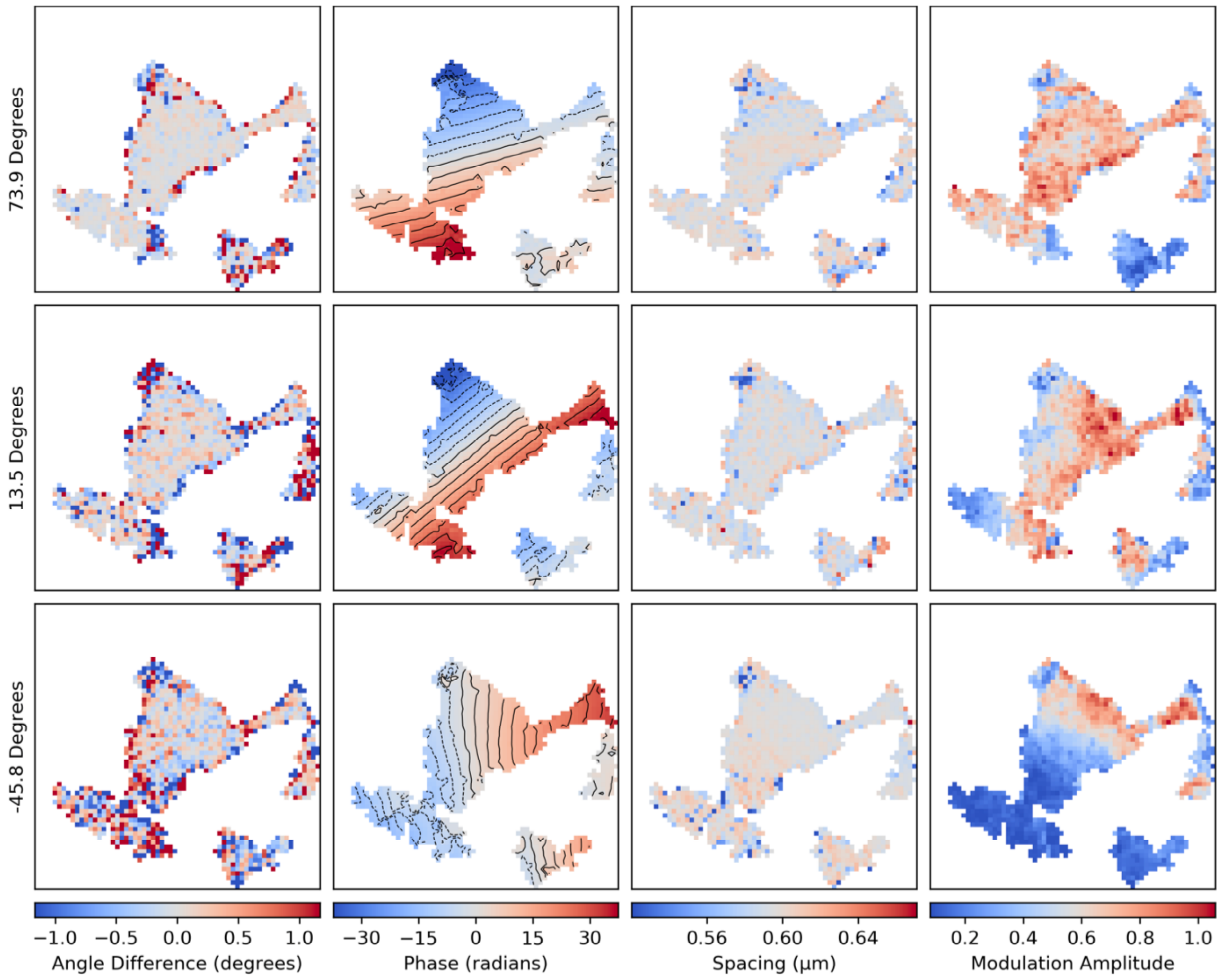


**Supplementary Fig. 2.** Maps of the spatial variation of illumination pattern parameters used in SR-SIM reconstruction.

Variability of illumination pattern parameters for each of the three orientations of illumination as measured during tiled SR-SIM reconstruction for the cell centered in the bottom of Fig. 1a.

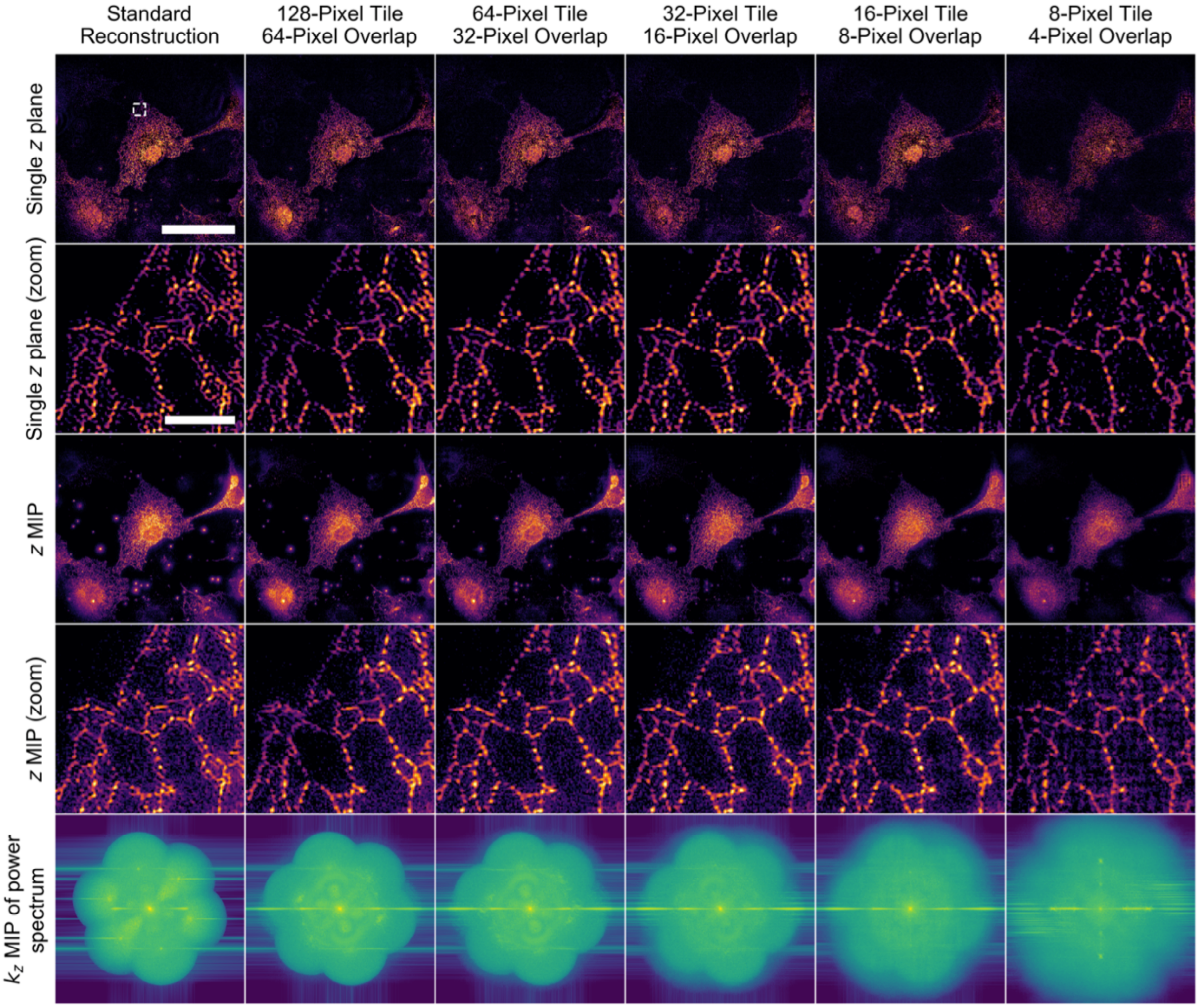


**Supplementary Fig. 3.** Effect of tile size on SR-SIM reconstruction.

Left column: Standard (i.e. no tiling) SR-SIM reconstruction. Remaining columns: tiled reconstructions using the same data, but with progressively smaller tile sizes. Real space data in columns 1-4 are shown with a gamma contrast adjustment of 0.5. Bottom row: xy MIP of the magnitude of the 3D Fourier transform for each of the reconstructed images, logarithmically scaled. White box: ROI displayed in the second and fourth rows. Even large tiles (e.g. column 2) can substantially improve image quality. Extremely small tiles, however, lead to pixilation artifacts in the final reconstruction as seen in both real (column 6, rows 1-4) and Fourier space (column 6, row 5). Tiled reconstruction is fairly insensitive to amount of tile overlap, but an overlap equal to half the tile width was chosen here as it did not lead to any noticeable stitching artifacts.

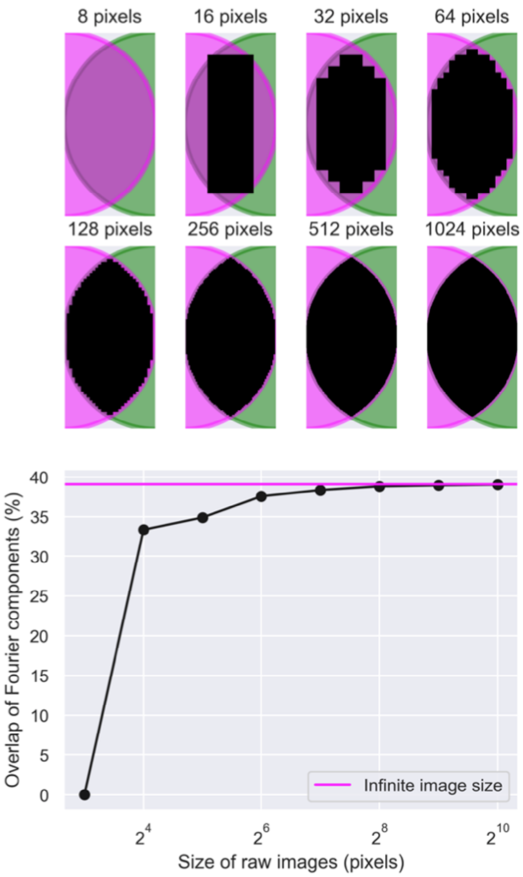


**Supplementary Fig. 4.** Effect of pixilation on the overlap of Fourier components for different SR-SIM reconstruction tile sizes**.**

A crucial step in the SR-SIM reconstruction process is determining the correct locations, in Fourier space, of the frequency-shifted components (e.g., magenta and green at top) that comprise the extended Fourier transform of the image. This is accomplished using an initial estimate based on the known illumination patterns, and then fine tuning the location based on the cross-correlation of spatial frequencies shared in their mutual overlap region (black at top) between neighboring transform components. At small tile sizes, there are an insufficient number of pixels in the overlap region, with too coarse a pixel size, to accurately fine tune the relative locations of the various components, reducing the accuracy of the SR-SIM reconstruction.
